# Insights into the Human Gut Virome by Sampling a Population from the Indian Subcontinent

**DOI:** 10.1101/2022.01.20.477175

**Authors:** Kanchan Bhardwaj, Anjali Garg, Abhay Deep Pandey, Himani Sharma, Manish Kumar, Sudhanshu Vrati

## Abstract

Gut virome plays an important role in human physiology but remains poorly understood. This study reports an investigation of the human gut DNA-virome of a previously unexplored ethnic population through metagenomics of faecal samples collected from individuals residing in Northern India. Analysis shows that, similar to the populations investigated earlier, majority of the identified virome belongs to bacteriophages and a smaller fraction (< 20%) consists of viruses that infect animals, archaea, protists, multiple domains or plants. However, crAss-like phages, in this population, are dominated by the genera VII, VIII and VI. Interestingly, it also reveals the presence of a virus family, *Sphaerolipoviridae,* which has not been detected in the human gut earlier. Viral families, *Siphoviridae*, *Myoviridae*, *Podoviridae*, *Microviridae*, *Herelleviridae* and *Phycodnaviridae* are detected in all of the analyzed individuals, which supports the existence of a core virome. Lysogeny-associated genes were found in less than 10% of the assembled genomes and a negative correlation was observed in the richness of bacterial and free-viral species, suggesting that the dominant lifestyle of gut phage is not lysogenic. This is in contrast to some of the earlier studies. Further, several hundred high-quality viral genomes were recovered. Detailed characterization of these genomes would be useful for understanding the biology of these viruses and their significance in human physiology.

**Importance:** Viruses are important constituents of the human gut microbiome but it remains poorly understood. The Indian subcontinent is a unique biogeographic region and the Indian population is known to harbour a distinct bacterial microbiome. However, the gut virome in this population has not been investigated earlier. Therefore, in this study, we investigated fecal samples of 12 healthy individuals to analyze their gut virome, through metagenomics.

## Introduction

The human gut harbours a microbial ecosystem that consists of trillions of microbes including bacteria, viruses, fungi, archaea and protists.^1^ Majority of the microbiome studies have focused on the bacterial component. Our understanding of the virome has been hampered, primarily due to technical challenges associated with analysis of viruses as well as due to limited availability of resources such as comprehensive viral databases and virus-specific tools for bioinformatic analyses.^2^ Nevertheless, metagenomics of faecal samples has provided significant insights into virome composition, led to discovery of novel viruses, revealed association with diseases and ethnic populations, and contributed to development of viral databases.^3–10^ Although, most of the data generated by metagenomics studies remains uncharacterized (86-99 % of the sequencing reads) and no consensus has emerged on the existence of a core gut virome, the identifiable fraction is reported to be dominated by DNA-bacteriophages in all studies.^7, 11^ Hence, in addition to understanding the viral composition, it has also been of interest to understand the dynamics between bacteriophages and bacteria in this ecosystem. Towards this, initial studies suggested that the human gut is dominated by temperate phages unlike other microbe-rich ecosystems such as the oceans.^12–15^ Although, a recent analysis by Shkoporov et al.^16^ has indicated that temperate bacteriophages do not dominate the human gut.^16^ Hence, further investigations are still required for establishing the nature of microbial dynamics in the gut as well as to address if there is existence of any core virome.

Several studies showing an association of gut virome with diseases as well as engraftment of phages in successful FMT trials have indicated the possibility of diagnostic and therapeutic use of gut phageome.^9, 10, 12, 18, 19^ However, for translational application of the virome, it would be important to establish the impact of various factors such as age, genetic background and geography on the variation of healthy virome.^16^

The Indian subcontinent represents a unique biogeographic region and the Indian population is known to harbour a distinct bacterial microbiome with respect to its diversity and the prevalent microbial taxa.^20^ It is possible that it will have a characteristic co-residing virome. Therefore, we investigated the gut virome by sampling a population from the Indian subcontinent. We analysed the DNA viruses present in the faecal samples of 12 “healthy” individuals by shotgun sequencing of the DNA isolated from purified virus-like-particles (VLPs) as well as from total microbial populations. Here, we present the composition of “healthy Indian gut virome” and demonstrate the presence of a virus family, *Sphaerolipoviridae,* which has not been previously reported in the human gut. We also demonstrate how the choice of sample preparation can impact virome analysis. Further, this study provides insights into the phylogenetic core gut virome and the viral lifestyle in the gut.

## Results

### (i) Effect of sample preparation on virome analysis

There are two methods for the isolation of viral DNA from fecal samples. One involves enrichment of virus-like-particles (VLPs) followed by extraction of the VLP-DNA. The second method involves extraction of total microbial DNA without enrichment for VLPs. In most of the earlier analyses of fecal viromes, one of the two methods has been used. There is only one recent study by Shokoporov et al. (2019) where both methods were used.^16^ Here, we also used both methods to extract DNA from each faecal sample to include free-viruses as well as host-associated-viruses in our analysis. Shotgun sequencing of the VLP-DNA and the total-microbial DNA generated 20-150 million reads per sample. The raw reads were filtered to remove low-quality sequences as well as any contaminating sequences of human origin (Figure 1A). We note that the recovery of high-quality sequences varied between samples although, equal starting material was used for each sample and the samples were processed similarly (Figure 1A).

**Figure 1.**
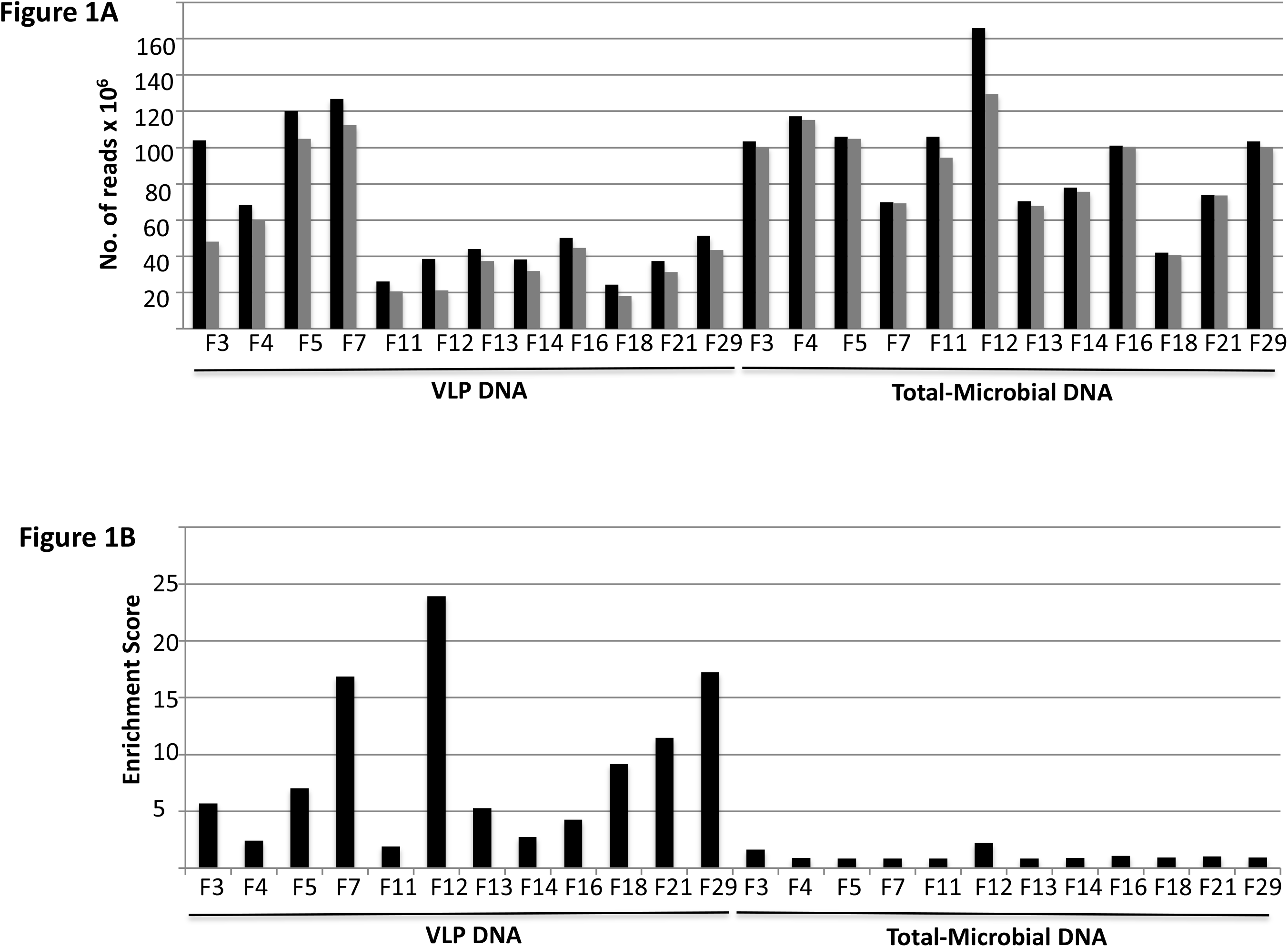
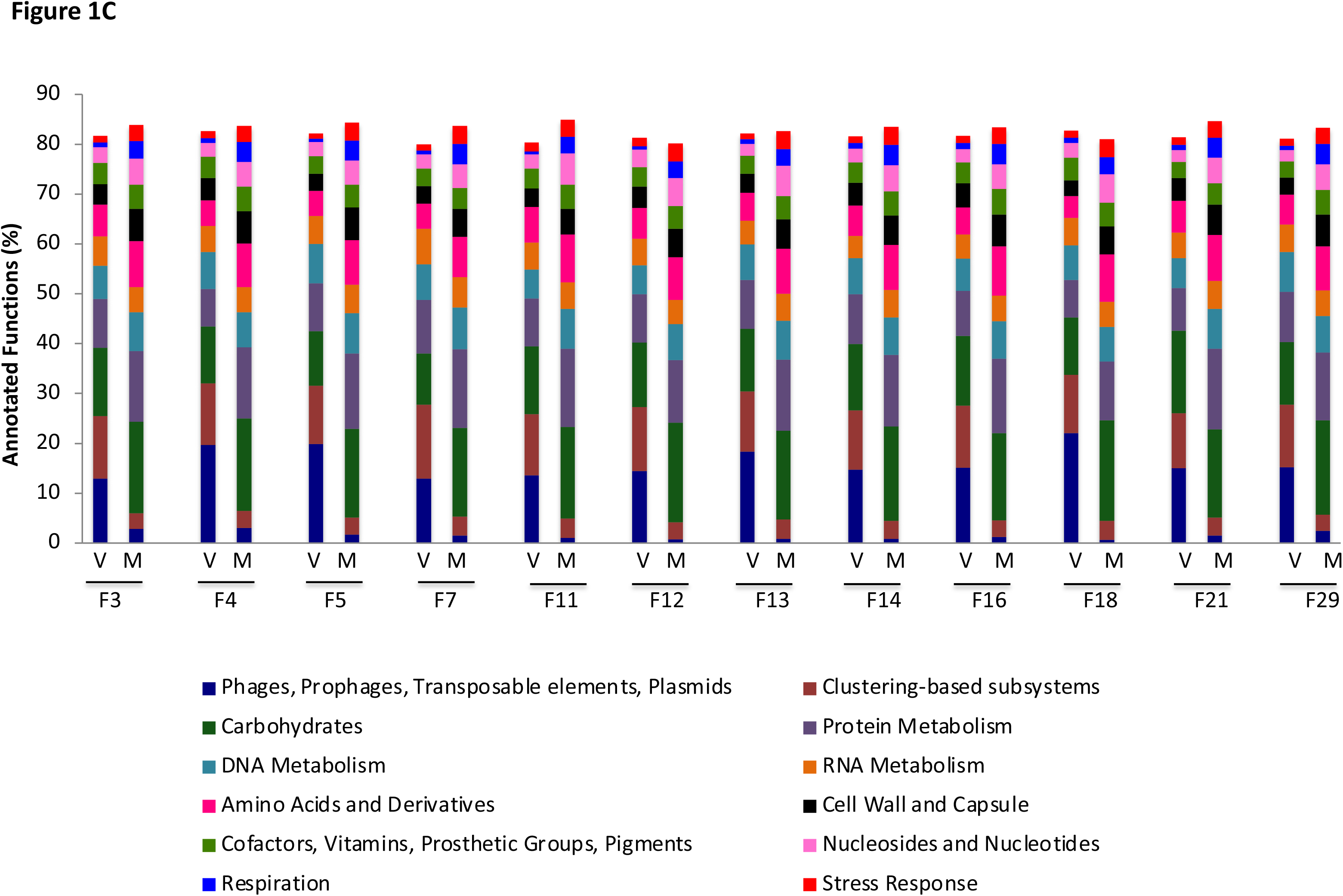
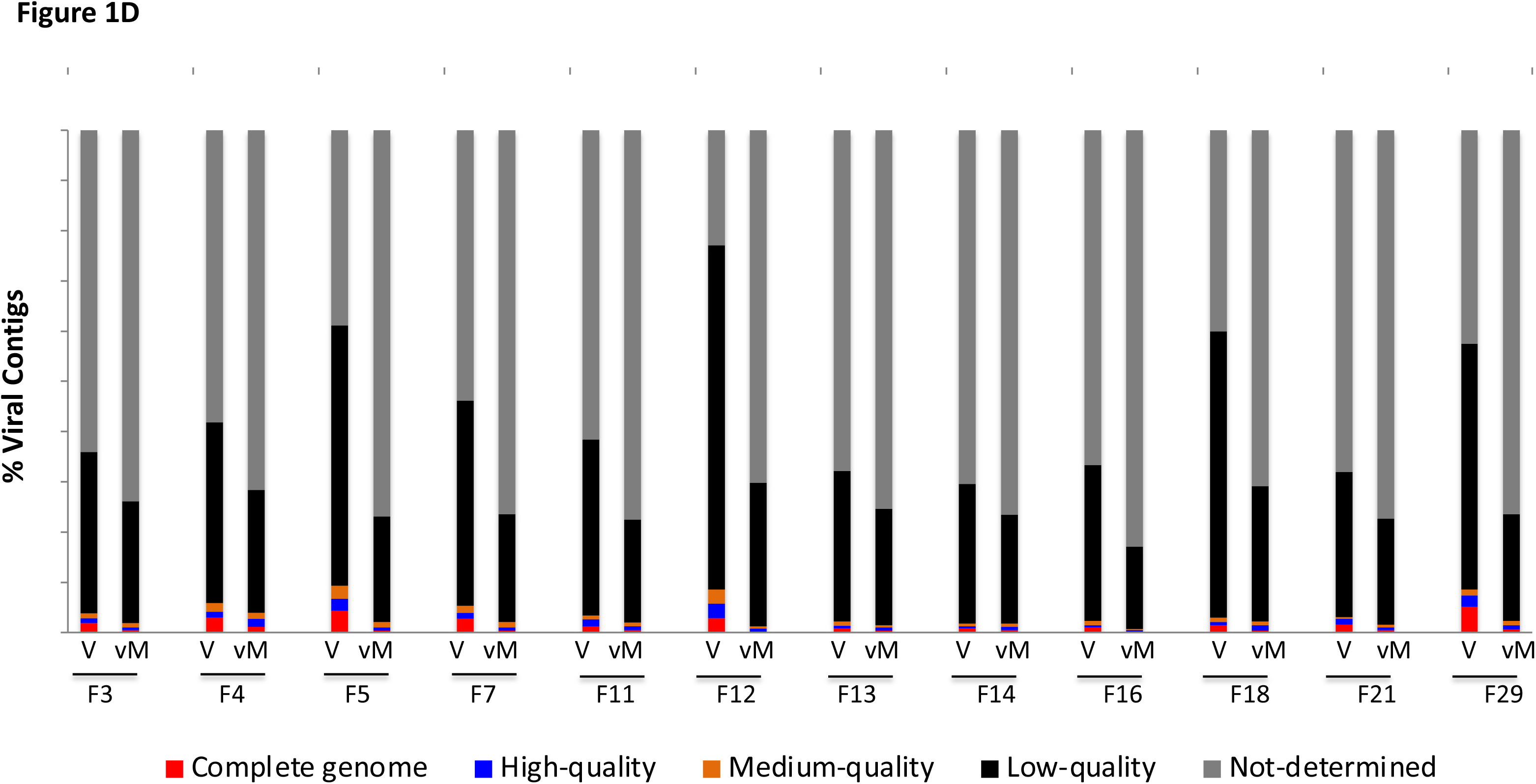
Data summary and quality check. (**A)** The number of reads before (black bars) and after (grey bars) quality control is shown. Subject identity and the type of samples are indicated on the x-axis. The VLP DNA represents both dsDNA and ssDNA obtained from purified VLPs. Total microbial DNA represents dsDNA. (**B)** Quality filtered reads were analyzed with ViromeQC tool. The y-axis represents the virus enrichment scores and the corresponding samples are indicated on the x-axis. (**C)** Functional profiles of the contigs obtained from the VLP-DNA and the total microbial DNA. Protein features were predicted and annotated by the MG-RAST pipeline. The abundance of the top most-abundant 12 categories in the Subsystems database is plotted for the samples identified on the x-axis. The identified classes are listed at the bottom of the graph and their proportion is indicated on the y-axis. (**D)** Mining tools were used to identify viral contigs after *de-novo* assembly and the redundancy was removed within each sample. The CheckV tool was used to analyze the quality of the assembled genomes. The proportion of viral contigs classified as “complete genomes”, “high-quality genomes”, “medium-quality genomes”, “low-quality” and “not-determined” in each sample are depicted. Subject identity is indicated on the x-axis. VLP-derived contigs are depicted as “V” and the total microbial DNA-derived viral contigs are depicted as “vM”.

Further, mapping of the quality-filtered reads to the NCBI viral RefSeq database showed alignment of 0.2 % of the VLP-DNA-derived and 0.05 % of the total microbial DNA reads (Table S1). Notably, only a limited fraction of the total reads could be mapped to the viral reference database, which is similar to what has been observed in earlier studies and has been attributed to the limited size of the reference databases. Nevertheless, the proportion of viral sequences in the VLP-DNA fractions was higher than in the total microbial DNA fractions (Table S1). Next, we analysed the read quality with the tool, ViromeQC. This tool is programmed to assign enrichment scores to sequences, based on their alignment to non-viral genes including small subunit and large subunit ribosomal RNA genes, and single-copy markers. Strikingly, enrichment scores assigned to the VLP-DNA-derived reads were significant (≥ 5.0) as well as higher than the scores assigned to the total microbial DNA-derived reads, indicating that the VLP fractions were significantly enriched for viral sequences and were free of contamination by the host DNA (Figure 1B). We again note that despite similar processing, there is variation among samples (Figure 1B).

Following metagenome assembly of the quality-filtered reads, we recovered around 6-fold more number of contigs (>1 kb) from the total microbial DNA fractions (5,90,939 contigs) as compared to the VLP-DNA fractions (93,612 contigs) (Table S2). This is possibly because in most samples we had recovered more number of high-quality raw reads from the total microbial DNA fractions as compared to the VLP-DNA fractions (Table S2). We analysed these contigs for their functional profiles, using the SEED subsystems database on the MG-RAST platform. Analysis revealed that the most abundant category of functions encoded by the VLP-DNA-derived contigs was “phages, prophages, transposable elements and plasmids” (13-20 %) followed by the “clustering-based subsystems” (12-15 %) (Figure 1C). The “clustering-based subsystems” represent those proteins for which the exact roles are not yet known. Whereas, functional profiles of the total microbial DNA-derived contigs showed a relatively higher proportion of genes related to “stress response”, “respiration”, “nucleosides and nucleotides”, “amino acids and derivatives” but significantly lower proportions of “clustering-based subsystems” (<4 %) and “phages, prophages, transposable elements and plasmids” functions (< 3 %) (Figure 1C). Detection of high levels of genes related to “phages, prophages, transposable elements and plasmids” further confirmed that VLP purification resulted in enrichment of viral sequences. This analysis also highlights that a significant proportion of viral genes encode for proteins with unknown functions.

For a more stringent selection of viral sequences from the total pool of contigs, we used three virus-mining tools, VirFinder, VirSorter and CAT. Notably, VirFinder identified a greater number of contigs as viral in the total microbial DNA fractions as compared to the VLP-DNA-derived fractions (Table S3). Whereas, VirSorter and CAT identified more number of contigs as viral in the VLP-DNA-derived fractions (Table S3). In total, around 35.7 % of the VLP-DNA-derived contigs (>1 kb) and around 7.3 % of the total microbial DNA-derived contigs (>1 kb) were identified as viral by the three tools (Table S3). The VLP-DNA- and the total-microbial DNA-derived viral contigs were pooled for individual samples and redundancy was removed within each sample. In total, we recovered 61,099 viral contigs (>1 kb), of which 36.6 % were derived from VLP DNA and 63.4% were from total microbial DNA (Table S3).

Next, we used the tool CheckV to examine the quality of the identified viral sequences. CheckV categorized the contigs as “complete genomes”, “high-quality” (more than 90 % complete genomes), “medium-quality” (50-90 % complete genomes), “low-quality” (0-50 % complete genomes) and “undetermined quality” (Figure 1D; Table S4). It determines completeness by comparing query sequences with the database of complete viral genomes followed by estimation of expected genome sizes. Genomes are categorized as “complete”, based on the presence of the direct terminal repeat (DTR) or inverted terminal repeat (ITR) sequences or provirus integration sites. Sequences are categorized as “undetermined quality” when they do not match any of the CheckV reference genomes and do not have any viral hidden Markov models (HMMs). Therefore, contigs that are classified as “undetermined quality” could represent very novel viruses, very short contigs or they may not be viral at all. In total, we recovered 600 “complete” viral genomes, out of which 437 were from VLP-DNA and 163 from total microbial DNA. Overall, these results show that viral sequences could be obtained with or without VLP enrichment. However, the sequences recovered by each method are unique. Further, the method of VLP enrichment before DNA extraction resulted in better recovery of “complete genomes”.

### (ii) Identification of known and unknown families of viruses

Read mapping to the NCBI viral RefSeq database identified the presence of bacteriophages (84.8 ± 17.6 %), animal viruses (4.29 ± 2.8 %), protist viruses (2.3 ± 3.17 %), archaeal viruses (0.11 ± 0.12 %), plant viruses (0.1 ± 0.2 %) and viruses that infect multiple domains (1.29 ± 1.1 %) (Figure 2A). Further, we annotated the viral contigs, either based on sequence alignment or by clustering of their predicted proteins. Protein alignment-based program, Kaiju, assigned taxonomic classification to ∼18 % of the total contigs (10,917 contigs out of the total 61,099) (Figure 2B). The 13 identified bacteriophage families (9,832 contigs), include *Siphoviridae*, *Myoviridae*, *Podoviridae*, *Herelleviridae*, *<u>Autographviridae</u>*, *Ackermannviridae*, *Microviridae*, *<u>Demerecviridae</u>*, *<u>Drexlerviridae</u>*, *Tectiviridae*, *<u>Chaseviridae</u>* and *Inoviridae*. The identified animal viruses (141 contigs) were from 13 families, *Herpesviridae*, *Iridoviridae*, *Ascoviridae*, *Poxviridae*, *Malacoherpesviridae*, *Circoviridae*, *Polyomaviridae*, *Alloherpesviridae*, *Asfarviridae*, *Adenoviridae*, *Lavidaviridae*, *Papillomaviridae* and *Parvoviridae*. Further, 518 contigs were assigned to a single-family *Phycodnaviridae*; 406 contigs as protist viruses of two families, *Mimiviridae* and *Marseilleviridae*; 17 Archaea viruses of two families*, Pithovirus sibericum* and *Sphaerolipoviridae*; 2 viruses of the *Genomoviridae* family, and 3 plant viruses of the *Caulimoviridae* family (Figure 2B). Similar to virome analyses in other ethnic groups, we note that the majority of the identifiable reads as well as the assembled genomes belong to bacteriophages. Although mapping to protozoan, invertebrate and plant viruses has been observed in earlier studies as well, their relationship with the human gut has not been proven.^21^ Additionally, it has been suggested that mapping to large dsDNA genomes of *Ascoviridae*, *Iridoviridae*, *Marseilleviridae*, *Mimiviridae*, *Nudiviridae*, *Phycodnaviridae*, *Pithovirus* and *Poxviridae* families could be due to misassignments.^21^ Therefore, we also used the protein clustering-based method for taxonomic annotation.

**Figure 2.**
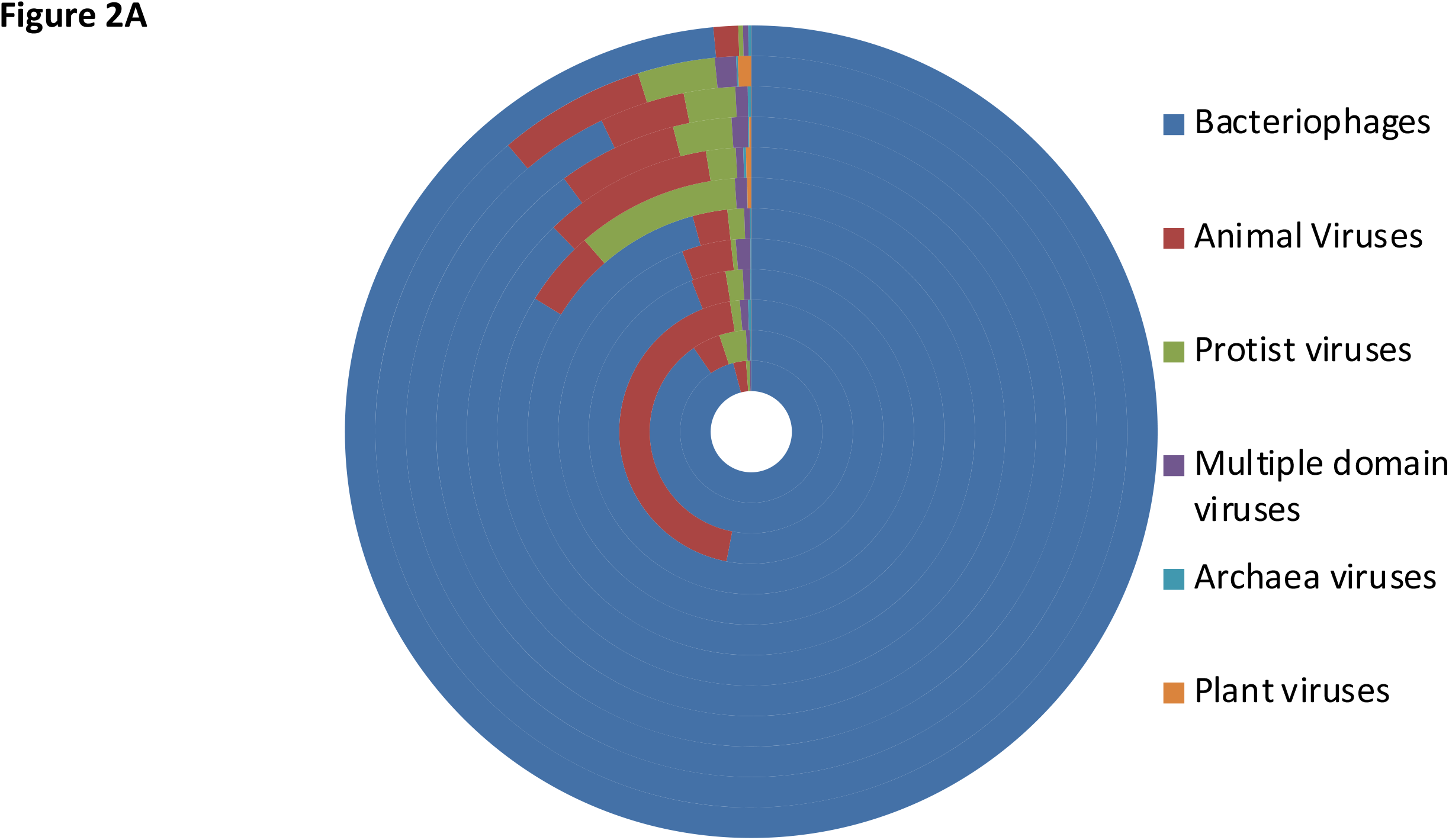
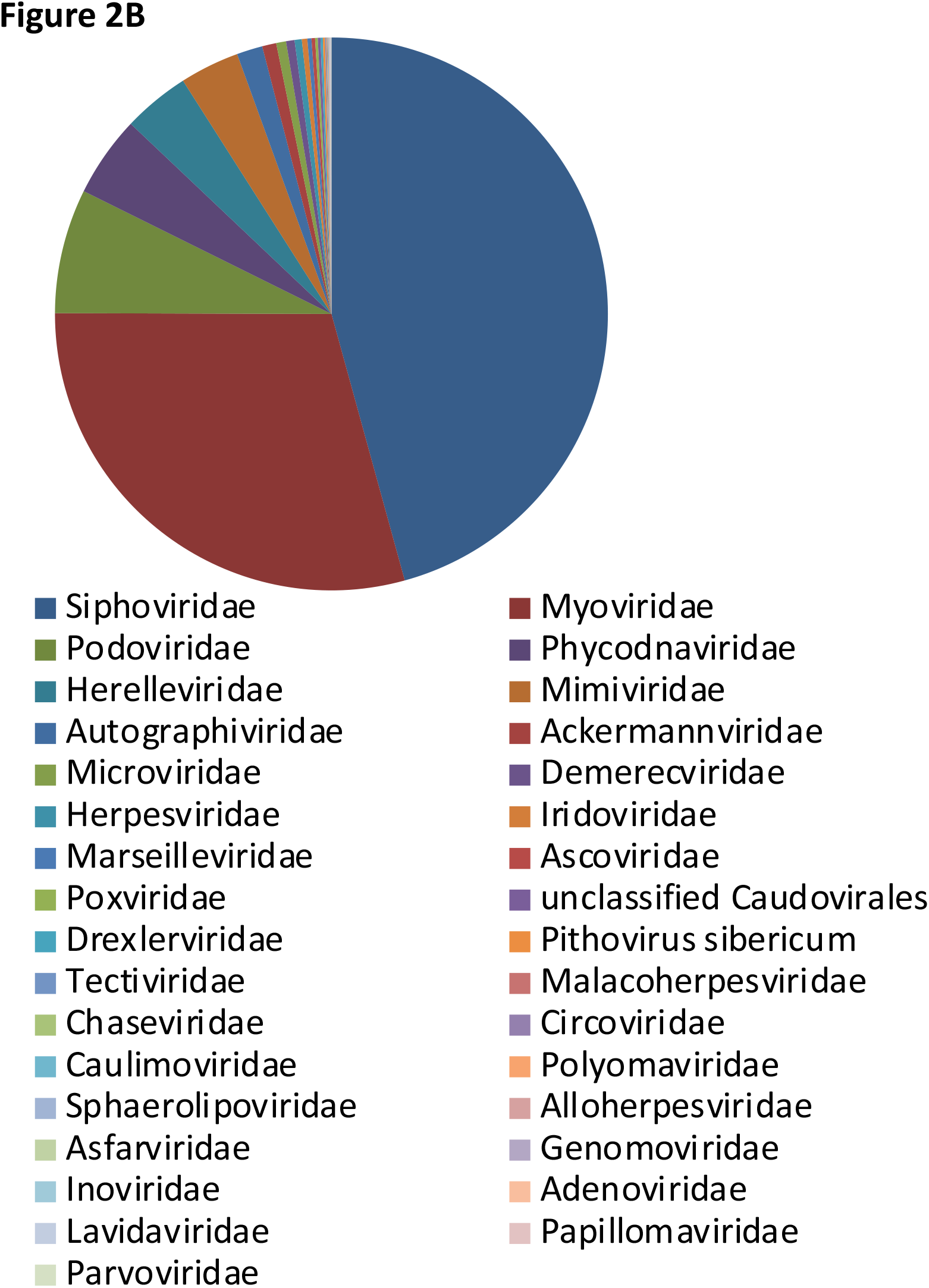
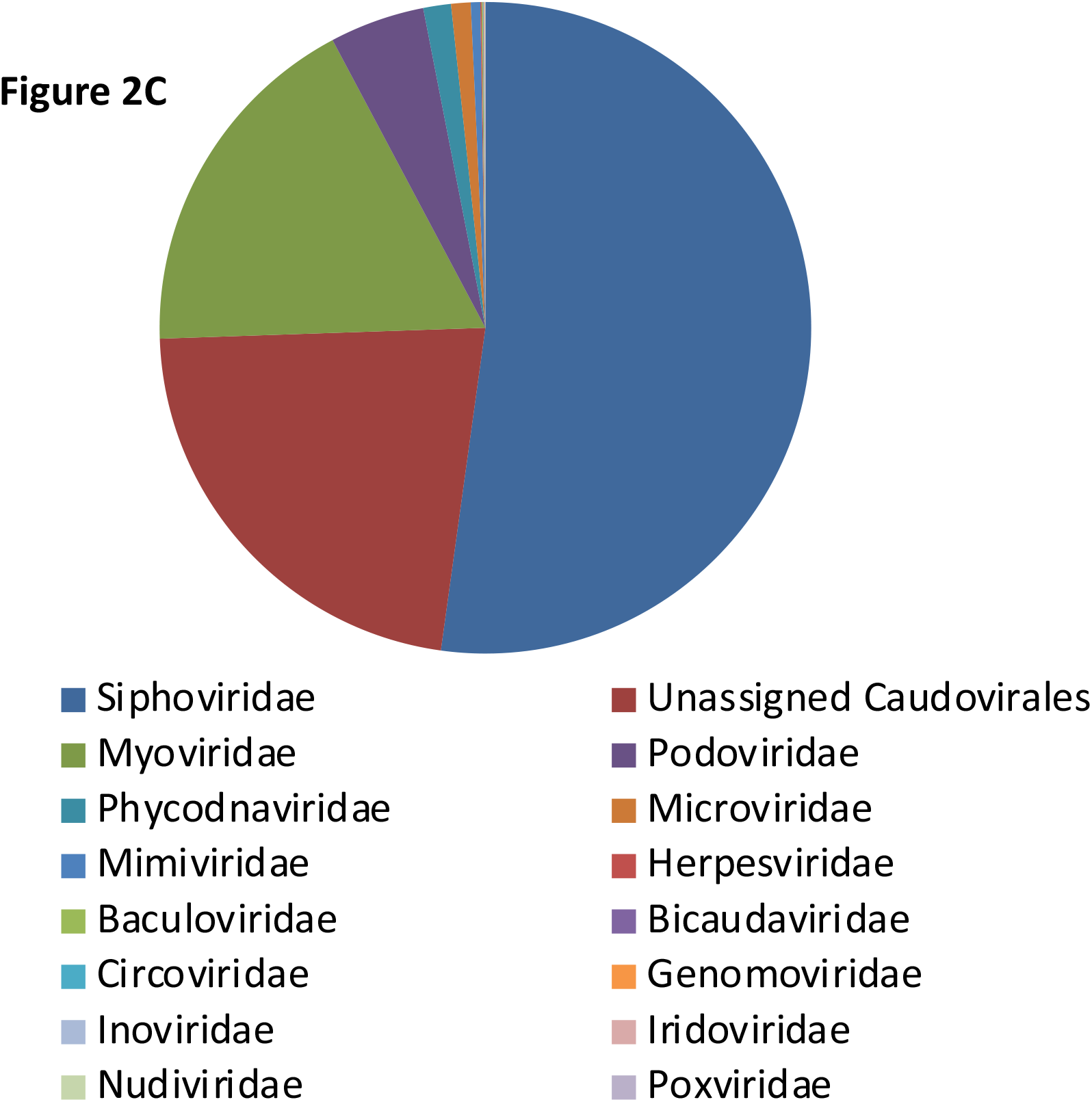
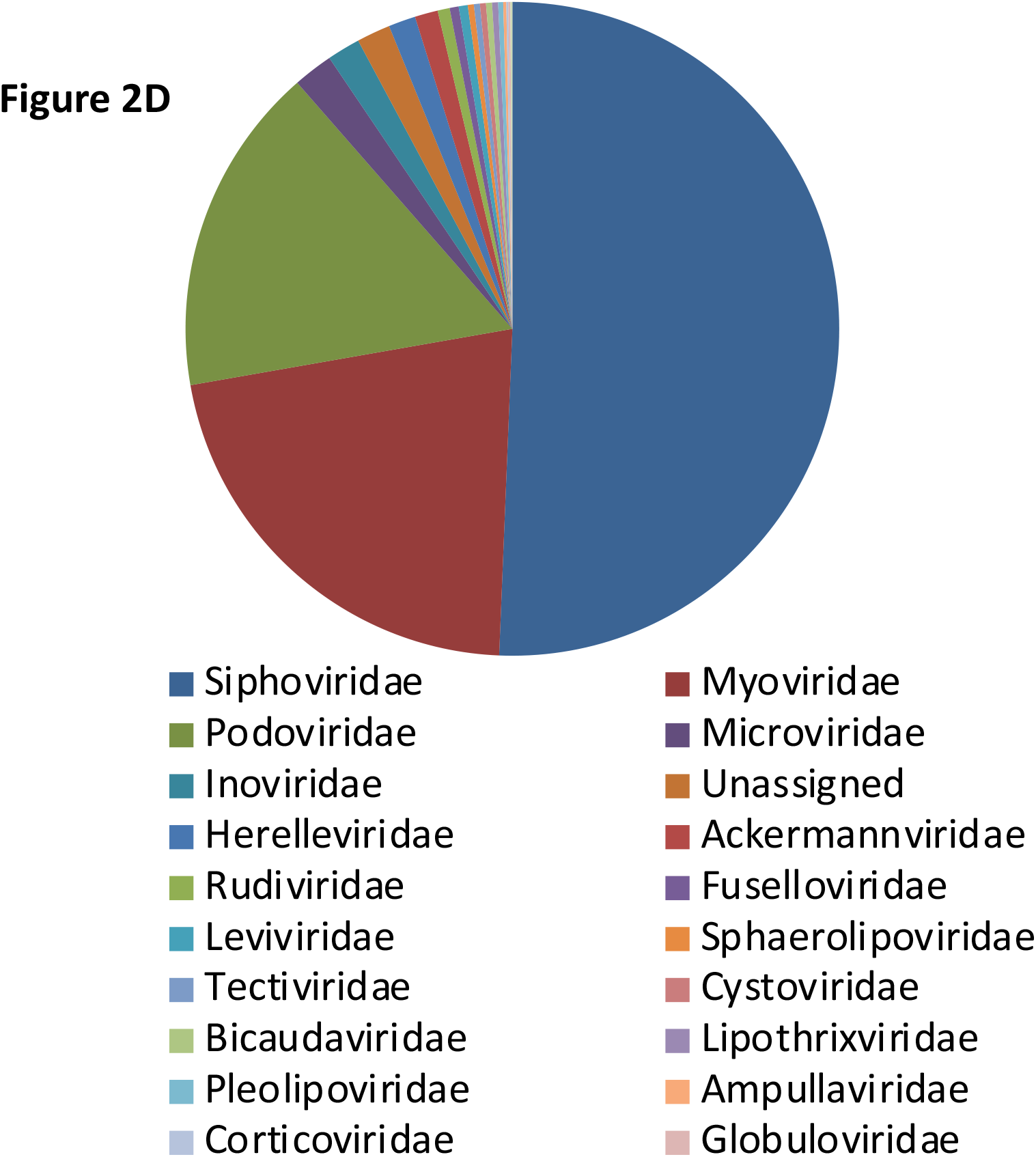
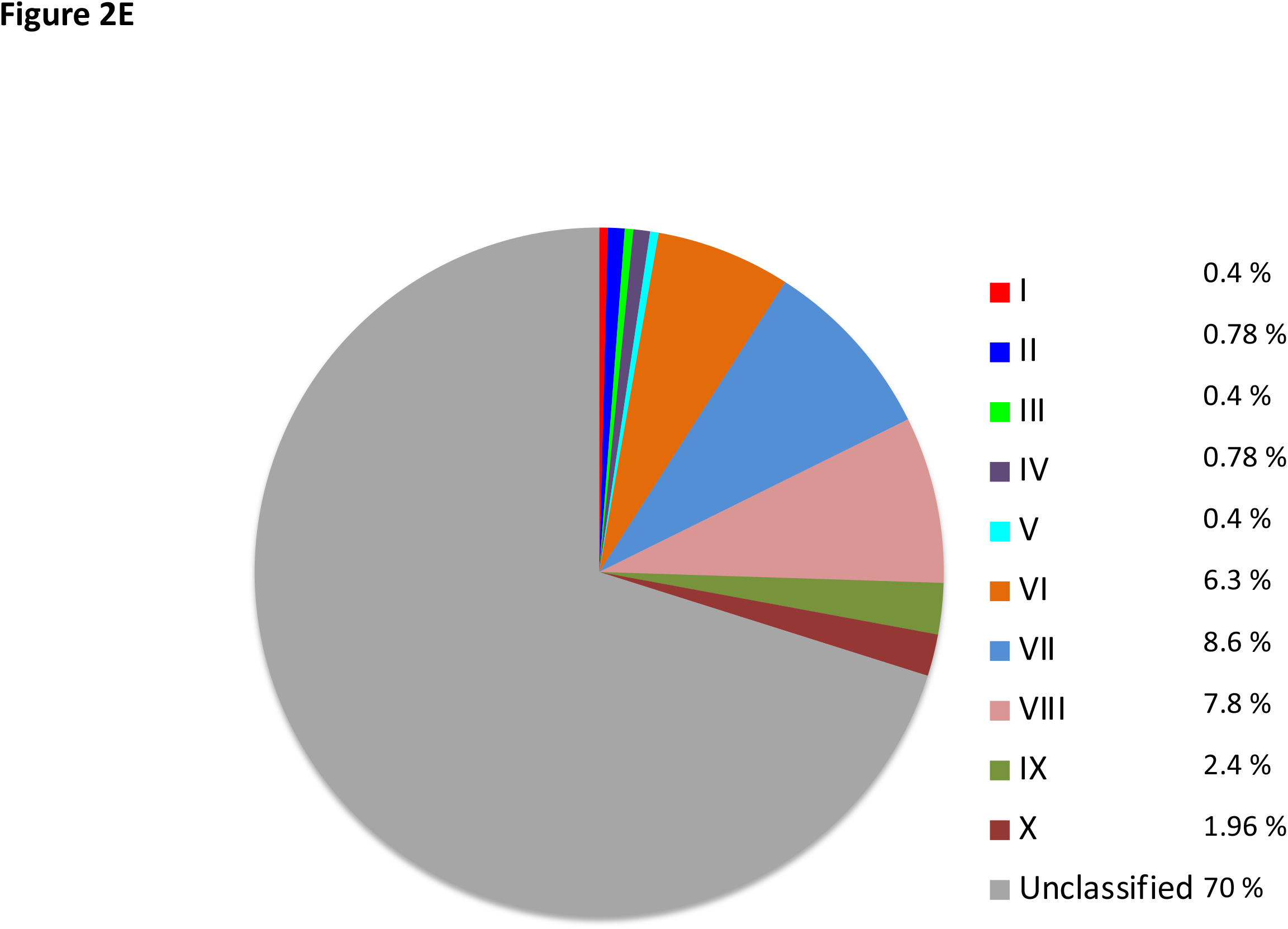
Taxonomic identification of viral sequences. (A) The high-quality reads mapped to the viral RefSeq database are depicted by a doughnut chart. Each of the viral categories is shown by a different color. Each ring shows the distribution of various viruses in a sample. The sequence of samples starting with the innermost ring and going outwards is F3, F4, F5, F7, F11, F12, F13, F14, F16, F18, F21 and F29. **(B)** Taxonomic assignment by the Kaiju program. Contigs that were identified as viral by the virus mining tools were assigned taxonomy based on protein alignment using Viral RefSeq databases, 1.1, 2.1, and 3.1. The pie chart is showing the identified viral families. **(C)** Taxonomic identification of the viral contigs based on the assignment of the vCONTACT2-generated clusters using Demovir and the TrEMBL database. The pie chart is showing the proportion of the identified viral families **(D)** Taxonomic identification of the viral contigs based on the assignment of the vCONTACT2-generated clusters using the ProkaryoticViralRefSeq94-Merged database. The pie chart is showing the proportion of the identified viral families. **(E)** The pie chart showing the proportion of each of the ten crAss-like phage genera and unclassified crAss-like phages.

Protein clustering was done with vConTACT2 and the clusters were identified with the help of two databases *i.e.* TrEMBL and ProkaryoticViralRefSeq94-Merged. Fifteen families were identified when the TrEMBL database was used for taxonomy assignment (Figure 2C). Except for *Baculoviridae*, *Bicaudaviridae* and *Nudiviridae,* all other families identified by this method were also identified by the protein-alignment-based method (Figure 2B). Among the animal viruses, *Herpesviridae* was detected by both methods. However, a significant proportion of the clusters remained unassigned (Figure 2C). The use of the ProkaryoticViralRefSeq94-Merged database resulted in the identification of 20 families of the prokaryotic viruses (Figure 2D).

Altogether, sixteen families including, *Siphoviridae*, *Myoviridae*, *Podoviridae*, *Herelleviridae*, *Microviridae, Inoviridae, Ackermannviridae, Phycodnaviridae, Mimiviridae, Sphaerolipoviridae, Tectiviridae, Circoviridae, Genomoviridae, Iridoviridae, Herpesviridae* and *Poxviridae* were identified by both, protein alignment as well as protein clustering-based methods (Figure 2B, 2C and 2D).

We also noted that the reads from all of our samples except one mapped to crAss-like phages (0.1-25 % of the total reads) (Table S5). Among the assembled metagenomes, we identified a total of 382 putative crAss-like phages (contigs >70 kb long), based on their sequence homology to seven most conserved proteins of crAss-like phages (Table S5). Around 30 % of these contigs could be classified into the ten known crAss-like phage genera, with VI, VII and VIII being the most dominant ones (Figure 2E). Further, around 43% of the identified crAss-like phages were found to contain at least one of the five lysogeny-related genes *i.e.* transposase, integrase, excisionase, resolvase and recombinase (Table S5).

### (iii) Evidence for the existence of a core virome

Although most of the viruses that are found in the human gut are unique to an individual, there are suggestions of a “core virome” or part of virome that is shared by individuals. However, its identity and the extent of sharing have varied in different studies, leaving the concept of a “core virome” debatable at this time.^12, 15, 16, 22^ Here, based on the protein alignment method, we found that 9 families out of 32 were present in all individuals (Figure 3A). These include *Siphoviridae*, *Myoviridae*, *Podoviridae*, *Phycodnaviridae*, *Herelleviridae, Mimiviridae, Microviridae, Demerecviridae and Herpesviridae*. When using the protein clustering-based method, the shared clusters identified with the TrEMBL database were *Siphoviridae*, *Myoviridae*, *Podoviridae*, *Microviridae* and *Phycodnaviridae* (Figure 3B). Whereas, the shared clusters identified with the ProkaryoticViralRefSeq94-Merged database belong to families *Siphoviridae*, *Myoviridae*, *Podoviridae*, *Microviridae*, *Inoviridae*, *Herelleviridae*, *Ackermannviridae*, *Rudiviridae*, *Fuselloviridae*, *Leviviridae*, *Sphaerolipoviridae*, *Tectiviridae*, *Cystoviridae*, *Bicaudaviridae*, *Lipothrixviridae, Pleolipoviridae*, *Ampullaviridae*, *Corticoviridae*, *Globuloviridae*, *Turriviridae* and *Phycodnaviridae* (Figure 3C). Collectively, the 6 families that were found shared among all individuals by both methods include *Siphoviridae*, *Myoviridae*, *Podoviridae*, *Microviridae*, *Herelleviridae,* and *Phycodnaviridae*.

**Figure 3.**
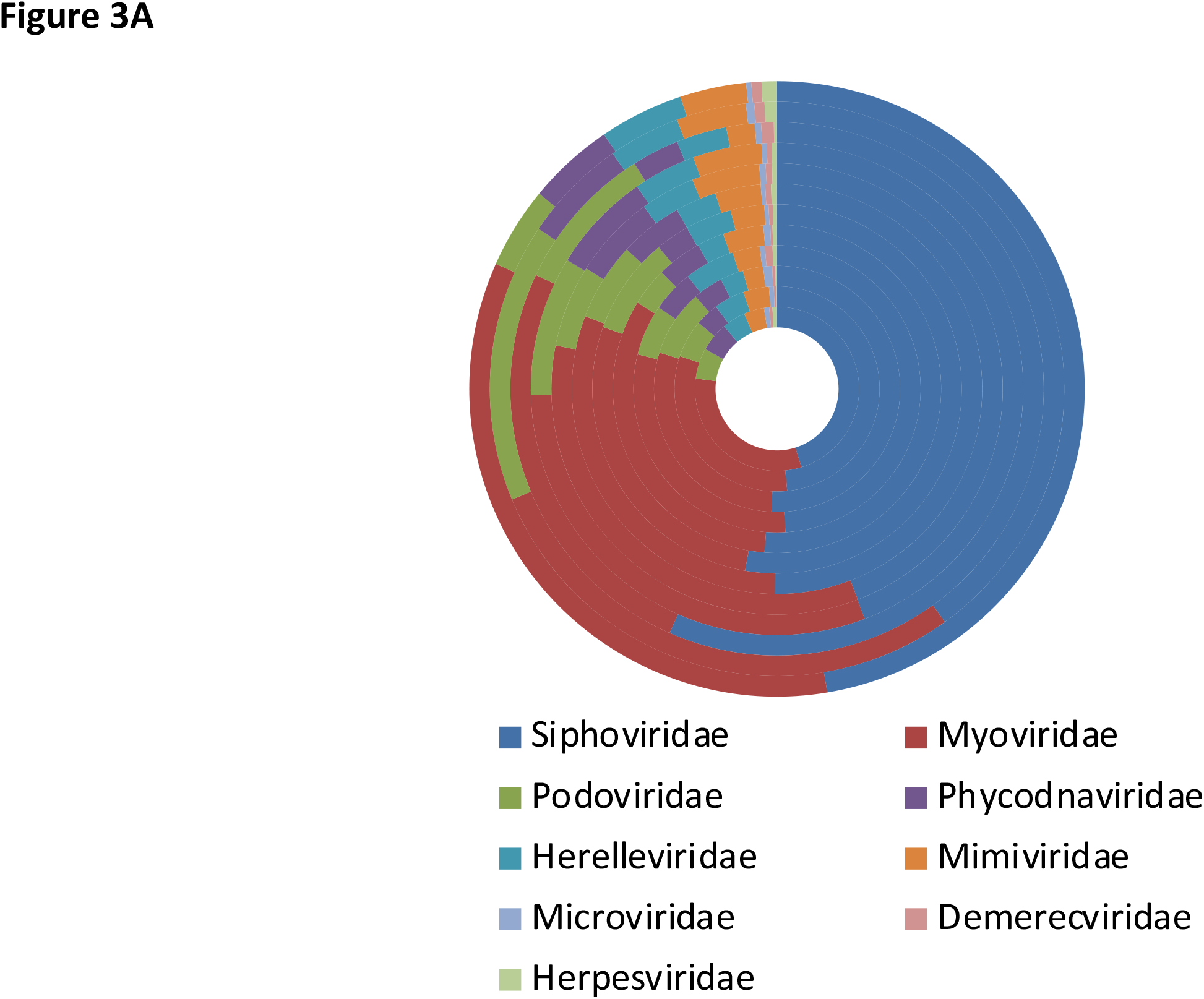
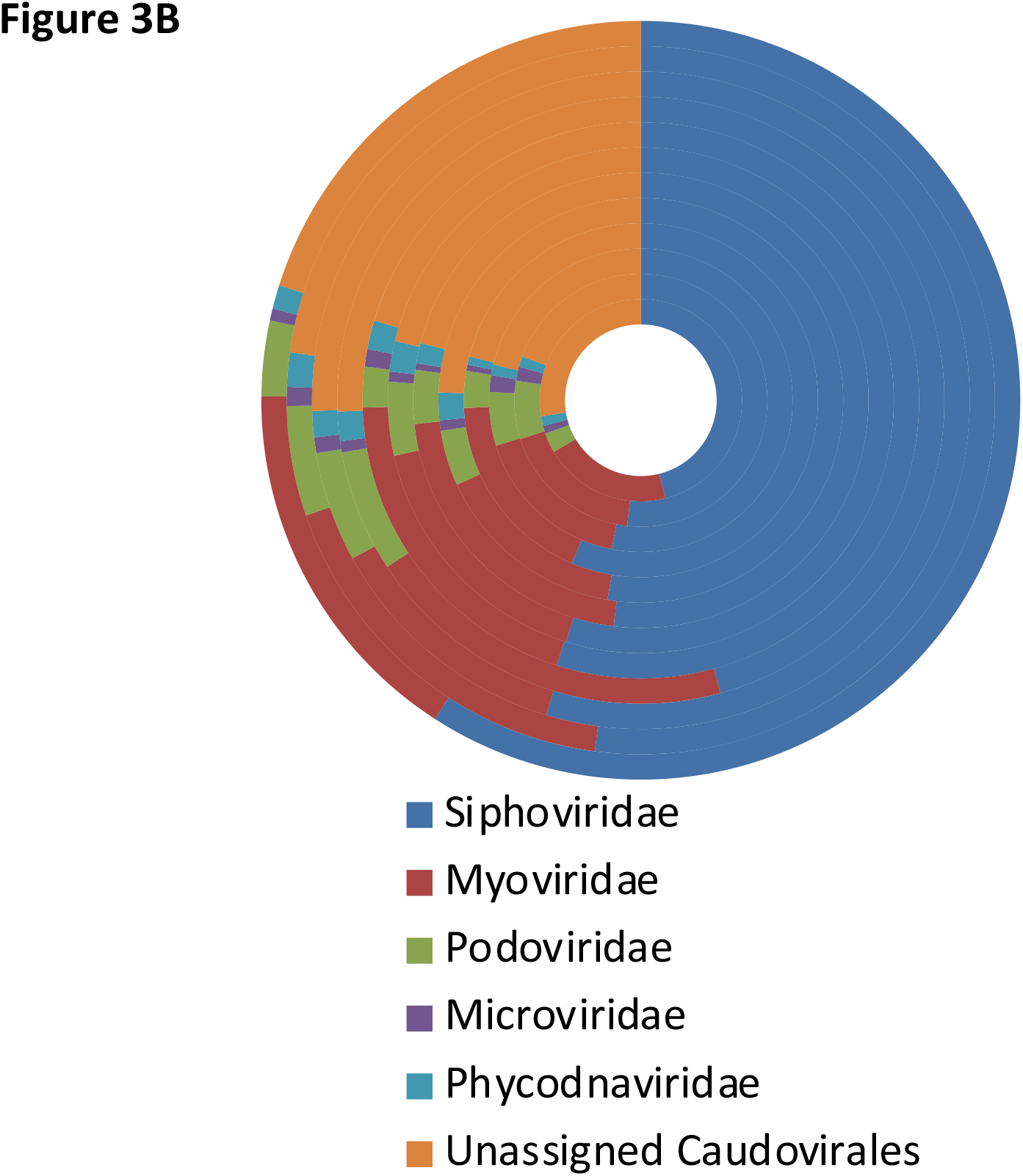
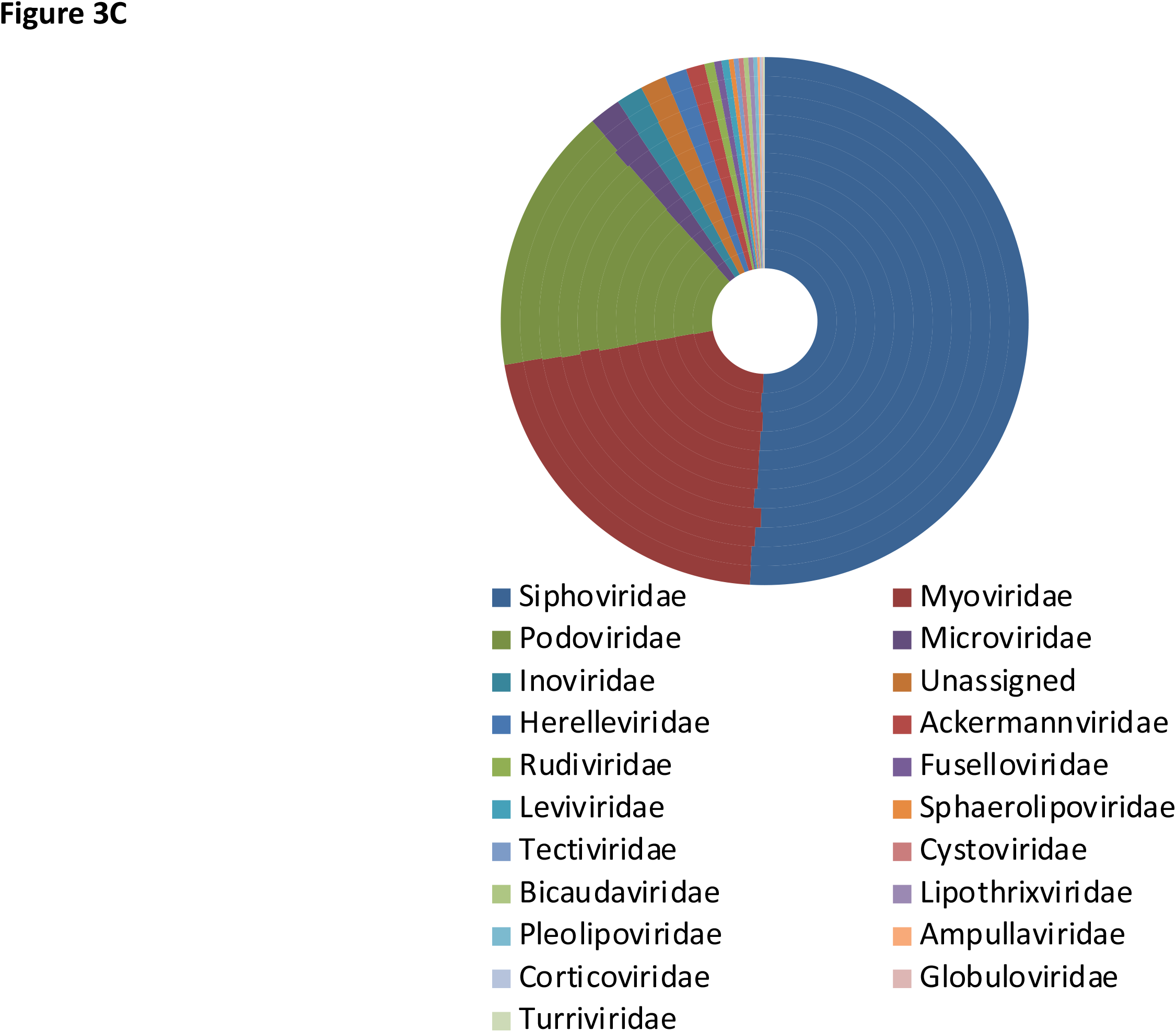
Identification of shared viral families. A doughnut chart showing viral families, that are shared across all samples. Each circle represents a sample. The sequence of samples starting with the innermost ring and going outwards is F3, F4, F5, F7, F11, F12, F13, F14, F16, F18, F21 and F29. **(A)** Viral families were identified by the protein-alignment-based method using the program Kaiju. Families that are present in all samples are shown. **(B)** Protein-based clustering of the viral contigs was performed by vConTACT2. Clusters were annotated using the TrEMBL database. Families shared by all samples are listed. **(C)** Clusters generated by vCONTACT2 were annotated using the ProkaryoticViralRefSeq94-Merged database. Families shared by all samples are listed.

### (iv) Lifestyles of the gut-resident bacteriophages and its effects on the co-residing bacterial population

The majority of the identified viruses in the gut are bacteriophages, which can significantly influence the structural and functional output of the ecosystem through processes such as host predation, lysogeny and horizontal gene transfer. Towards understanding their interaction with the co-residing bacterial population, we determined the richness of the bacterial as well as the viral species derived either from VLP-DNA or from total microbial DNA fractions (Figure 4A). The analysis revealed a moderate negative correlation between the richness of bacterial species and the VLP-DNA-derived viral species (r = −0.6; p = 0.037) but no significant correlation was observed with the total microbial DNA-derived viral species (r = −0.12; p = 0.71) (Figure 4B). Generally, VLPs (free virus particles) represent those viruses, which are undergoing lytic cycles. Whereas, total microbial DNA-associated viruses represent those, which are in a lysogenic state with the host; either integrated into the host genomes or existing as an extrachromosomal entity such as plasmids. Our results showing a negative correlation between bacterial and free viral (lytic) species richness align with the idea that dominance by phages that are undergoing lytic lifecycle results in reduced bacterial abundance. Further, we observed no correlation between the richness of bacterial species and the total microbial DNA-derived viral species (lysogenic). This is an unexpected result because an increase in the abundance of the bacterial population would be expected to increase the abundance of lysogenic phages. However, no correlation is possible if the number of lysogens present in the analysed population were low. Therefore, we determined the proportion of lysogenic viruses in our population by scanning our vOTUs with HMM profiles of five lysogeny-associated genes (transposase, integrase, excisionase, resolvase and recombinase). The vOTUs were assigned lysogenic if at least one of these lysogeny-associated genes was present in them. Across all individuals, we found that less than 10 % of the vOTUs contained lysogeny-associated genes (Figure 5A). Further, majority of the lysogenic vOTUs were derived from the total microbial DNA fractions (40-97 %) followed by the VLP fractions (3-57 %) and some were represented in both fractions (0-11%) (Figure 5B).

**Figure 4.**
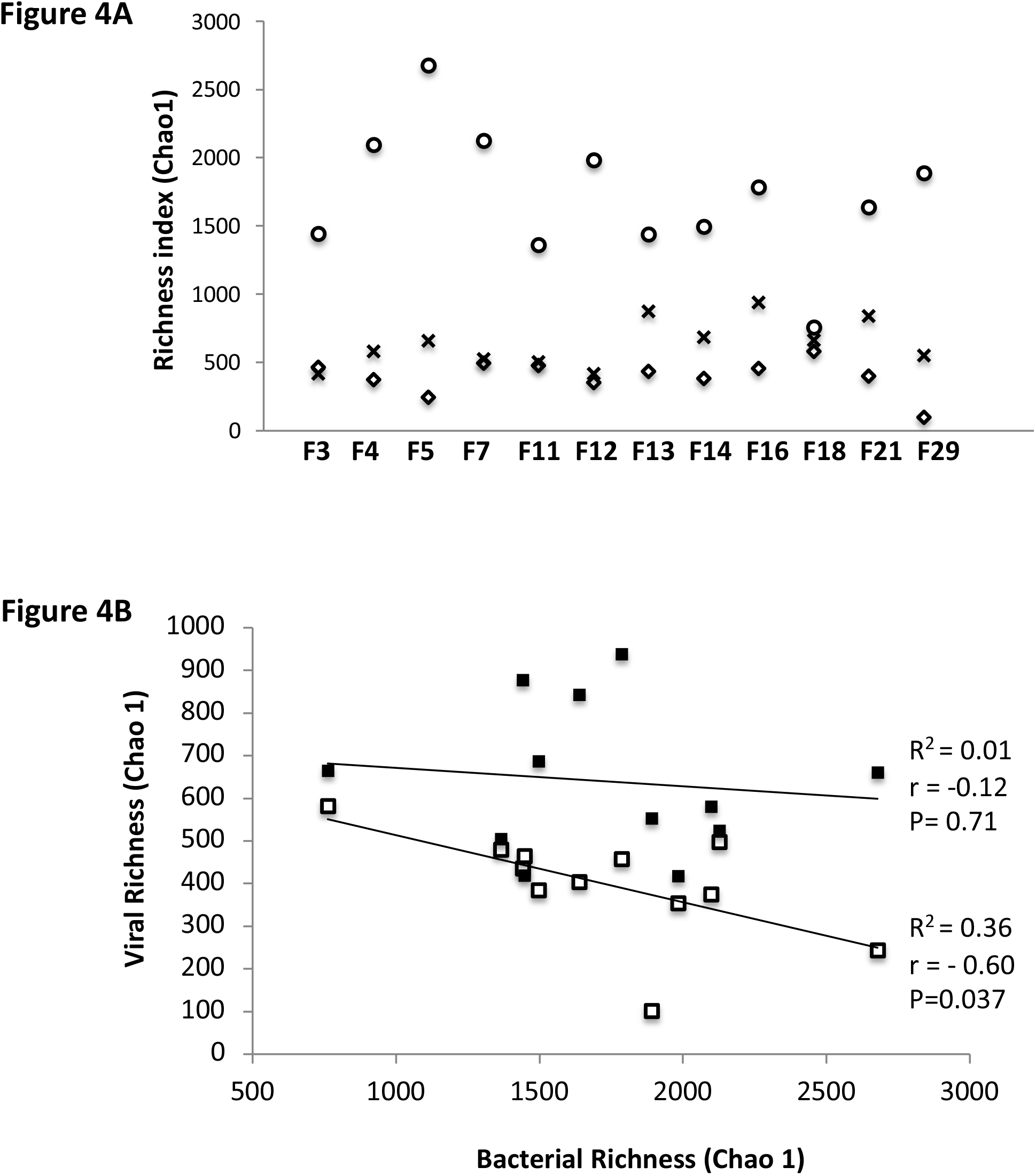
Diversities of the viral and bacterial populations and correlation analyses. (**A)** Species richness was determined with the Vegan package of R. Chao 1 indices are depicted on the y-axis with respective samples on the x-axis. The richness of bacterial species, phages that are associated with the host and the free virus (VLP) are represented with black bars, gray bars and white bars, respectively. (**B)** Species richness correlations of bacterial species richness with free viral species (solid line and filled circles; coefficient of correlation (r) is –0.12) as well as with phages associated with hosts (dotted line and open circles; coefficient of correlation (r) is – 0.60) are shown.

**Figure 5.**
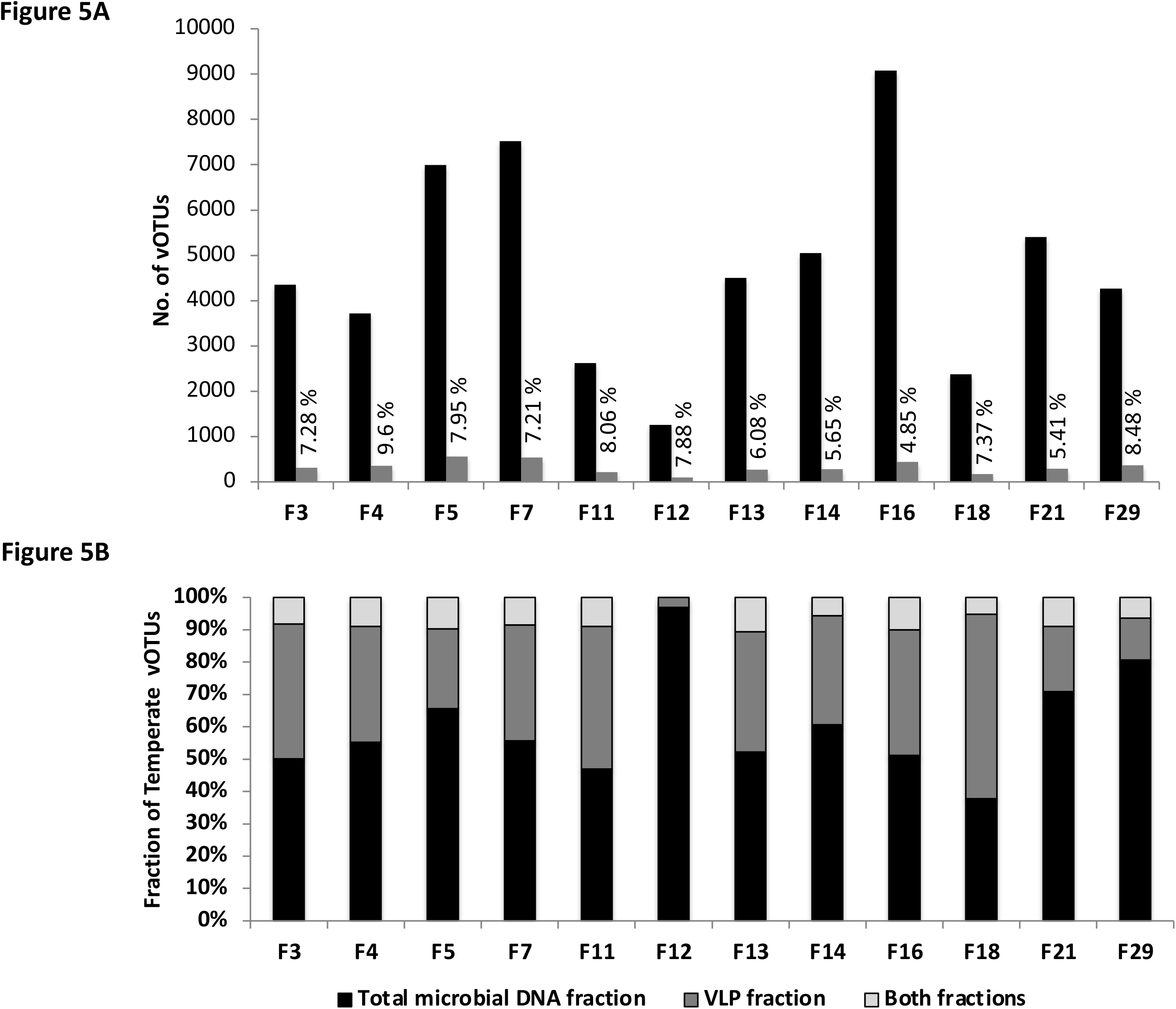
Bacteriophage lifestyle. Viral contigs were clustered as vOTUs with ClusterGenomes and ORFs were predicted in each vOTU, using Prodigal. ORFs were annotated using a custom set of HMM profiles of lysogeny-associated genes. **(A)** Black bars show the total number of vOTUs (y-axis) in each sample (x-axis) and the grey bars show the number of vOTUs in which lysogeny-associated genes were detected. The calculated percentage of lysogenic vOTUs is indicated on top of the bars. **(B)** The origin of the identified lysogenic vOTUs is shown.

## Discussion

The process of sample preparation for virome analysis is a significant consideration. In most of the previous metagenome analyses of human gut virome, samples had been prepared either by the enrichment of the VLPs before DNA extraction or by extraction of the total microbial DNA without enrichment. There is only one study where both methods were used to analyse 10 samples. Extraction of the total microbial DNA without enrichment is attractive and could be preferred for multiple reasons such as it provides convenience, is more economical, faster and can be adapted for simultaneous processing of a large number of samples. However, it is important to establish the effects of sample preparation. To address this, Gregory et al.^7^ performed bioinformatic analysis of the available metagenome data, collected in different gut virome studies. Their analysis revealed that (i) there is no significant effect of the two methods on the number of viral contigs assembled per bp sequenced; (ii) sample preparation method also showed no effect on the contig length, when data from different studies were analysed but longer contigs were detected with VLP-DNA-derived metagenomes when data from a single study which used both methods, were compared; (iii) the rate of viral detection was higher with the data obtained from the total microbial DNA extraction method; and (iv) the two methods capture different subsets of viruses.^7^ Similarly, our results also show that each method recovers a comparable number of viral contigs and that the viral sequences captured by the two methods are unique. In addition, we found that the recovery of “complete viral genomes” is better with the VLP enrichment method. This could be because the VLP-DNA fractions are significantly enriched with sequences of viral origin, which possibly led to better genome assembly. Our analysis also shows that the total microbial DNA fraction is enriched with phages containing lysogeny-associated genes, although not all. Therefore, data obtained through the total microbial DNA method could be useful for understanding the mechanisms related to the co-existence of phages with their hosts. However, concurrent analysis by both methods would be needed for a comprehensive understanding of a virome.

Taxonomic assignment of our data reveals that the gut virome is dominated by bacteriophages, which is similar to what has been observed in other populations. Although, one of the viral families, *Sphaerolipoviridae*, found in our samples has not been reported in the human gut earlier. Bacteriophages of the *Herelleviridae* family have been reported in one of the recent studies involving cohorts of Chinese and migrant Pakistani populations.^24^ Strikingly, we did not detect viruses of the *Anelloviridae* family, which are the predominant animal viruses in the Western population.^7^ The most abundant animal virus family in our samples was *Herpesviridae*. Based on the analysis by multiple methods, we found that members of at least six families, *Siphoviridae*, *Myoviridae*, *Podoviridae*, *Microviridae*, *Herelleviridae,* and *Phycodnaviridae* were present in all individuals. Since this analysis is based on a small fraction of identifiable sequences of the whole virome, it is not possible to estimate the extent of sharing between individuals. However, it does provide support for the existence of a phylogenetic core virome. Families of the order Caudovirales (*Siphoviridae*, *Myoviridae*, *Podoviridae*) and *Microviridae* have been known to form a phylogenetic core phageome.^3, 15, 22^ Interestingly, we found that in addition to these phage families, members of the *Herelleviridae* and *Phycodnaviridae* families were present at significant levels and were shared among all individuals. *Herelleviridae* is a relatively new family of phages added to the order Caudovirales. Viruses of the *Phycodnaviridae* family are large dsDNA viruses that are known to infect algae and there are pieces of evidence that they can infect humans as well.^25^ Further, we detected crAss-like phages in all except one sample. Since the discovery of crAssphage in 2018, the family of crAss-like phages has gained special interest because they are known as the most abundant phages in the human gut and are quite ubiquitous as well.^26^ In some of the gut viromes, up to 90% of the sequences are comprised of crAss-like phages. They have been classified into 4 subfamilies and 10 genera.^23, 27^ Using similar methods, we were able to classify only 30 % of our crAss-like phages, with genera VI, VII and VIII being the most represented. In the Western population, the most common genus is I and in the Malawian cohorts, they are genera VIII and IX. The significant fraction that remains unclassified requires further investigation.

Bacteriophages are important components of the gut ecosystem and therefore, understanding the mechanism(s) of phageome maintenance in the gut has been of interest. Apart from environmental conditions of the gut as well as the host-defense and phage counter defense system, phage lifestyle plays a significant role in this. However, consensus about the phage lifestyle in the human gut has not been reached. Our results showed a negative correlation in the richness of bacterial species and the VLP-derived viral species. VLP-derived viruses represent those viruses, which are produced upon the lytic cycle. These results, therefore, demonstrate the effect of the lytic lifestyle of phages on the co-residing bacterial population in the gut. Further, we detected lysogeny-associated genes only in a small fraction of viral sequences (<10 %). These results further suggest that the lysogenic lifestyle does not dominate in the gut. Many of the earlier investigations of gut virome have suggested that lysogenic lifestyle dominates in the gut.^12, 15, 17^ However, our results corroborate with one of the recent studies, which also reported that temperate phages do not dominate the gut ecosystem.^16^ Similarly, although crAss-like phages were initially predicted to have temperate lifestyle, evidences are emerging to suggest the existence of alternative lifestyles.^23^ In our analysis, we found lysogeny-associated genes in 43 % of the crAss-like phages. All these results suggest that in addition to lytic and lysogenic lifestyles, alternative phage lifestyles such as pseudolysogeny, chronic infection and carrier state might be operative for the maintenance of phages and their hosts in the gut ecosystem. However, further investigations will be needed to fully understand their existence and contribution.

This study has generated data on the virome comprising only the DNA containing viruses. To produce a complete picture of the “healthy Indian gut virome” attempts are underway to identify the RNA-genome containing resident viruses of the gut. The recovered “complete genomes” are also being analyzed further according to the guidelines provided by the Genomic Standards Consortium for reporting the sequences of uncultivated viral genomes (UViG).

## Methods

### 1. Subjec recruitment and sample collection

We obtained ethics clearances from the Institutional Biosafety and Human Ethics Committees (Reference No. RCB-IEC-H-14), recruited pre-defined “healthy” individuals, and collected faecal samples from them after obtaining written consent. Individuals between the ages of 20 and 35 years, who had normal body mass index, normal bowel frequency, no history of chronic intestinal disease or autoimmunity, had balanced meals at regular intervals, and had not received antibiotics in the 6 months before sampling were defined as “healthy”. Samples were collected using sterile containers, placed on ice immediately after collection, and then stored in a deep freezer, to be used within 3 months of collection.

### 2. DNA extraction and sequencing

VLP purification was done by sequential centrifugation, filtration and gradient ultracentrifugation. Homogenized extracts of the faecal samples were prepared by dilution of specimens in ice-cold SM (sodium and magnesium) buffer followed by thorough vortexing in presence of glass beads. Extracts were centrifuged at 4500g for 30 min at 4 °C, to remove particulate material. The supernatants were filtered sequentially through 0.45 μm and 0.22 μm-pore-size membranes. The filtrates were centrifuged on a step gradient of iodixanol. Before isolation of DNA from the purified VLPs, we treated them with DNase I and Benzonase to remove any free nucleic acid of human and bacterial origin that may co-purify during our procedure. To confirm the removal of contaminating bacterial DNA, VLP preparations were screened by 16S rDNA PCR. DNA was extracted from those VLP preparations that resulted in no amplification of DNA fragments with primers targeted for 16S rDNA (data not shown). VLP-DNA was extracted using the phenol:chloroform method. For the extraction of total microbial DNA, homogenized extracts of 200 mg fecal specimens were treated with enzymes, detergent and guanidine thiocyanate to lyse microbial cell walls and membranes followed by mechanical disruption using a bead beater. Insoluble fractions were removed by centrifugation and the total microbial DNA was isolated from the supernatants through conventional methods of nucleic acid precipitation. RNA was removed by digestion with RNaseA.^28^ Purified DNA was quantitated and its quality was checked before sequencing. Whole-genome shotgun sequencing was performed for all samples. Whole-genome metagenome sequencing libraries were prepared using the TruSeq DNA PCR-Free library preparation kit. Briefly, DNA was sheared using Covaris ultra sonicator. The fragmented DNA was end-repaired to remove any overhangs resulting from sonication. Library size was selected using sample purification beads. The size-selected molecules were mono-adenylated at the 3’-end followed by ligation of Illumina indexed adapters. The adapter-ligated fragments were cleaned up using purification beads and the clean fragments were assessed for size distribution on Agilent TapeStation. To include ssDNA viruses in the analysis, VLP DNA was PCR-amplified before shotgun sequencing, using Swiftbio kit from Illumina. In brief, whole-genome metagenome sequencing libraries were prepared using Accel-NGS 1S Plus DNA Library Kit, which processes both ssDNA and dsDNA in a mixed sample type. DNA was sheared using Covaris ultra sonicator and then denatured for adaptase step wherein 3’ tailing and ligation of truncated adapter 1 takes place. Next, the second strand of DNA is generated followed by truncated adapter 2 ligation in an extension and ligation step. The fragments were indexed using a limited cycle PCR, cleaned up using purification beads, and assessed for size distribution using Agilent TapeStation. The resulting libraries were quantitated and loaded on the cBot for cluster generation. Sequencing was performed on Illumina HiSeq X Ten platform with a read length of 150×2 bp.

### 3. Pre-processing of the sequencing data and metagenome assembly

The raw reads were processed using fastq-mcf (v1.1.2) to ensure that the data do not contain sequencing artifacts (Q_30_), sequence duplication and adapter sequences. Reads with an average quality Phred score below 30, low-quality tails in the reads and reads shorter than 36 bases were eliminated. Further, the high-quality reads were also filtered for human DNA contamination by aligning the reads to the human reference genome (hg19/GRCh37) using the Burrows-Wheeler aligner (BWA) and only the unaligned reads were taken for further processing. Filtered total microbial DNA reads were then aligned to bacterial, fungal, viral and archaea genomes in the NCBI RefSeq database. To evaluate the purity of the VLP samples, ViromeQC (v1.0.1) was used.^29^ Metagenome assembly was performed with MEGAHIT (v 1.2.9).

### 4. Identification of viral sequences

Contigs from MEGAHIT were filtered for size >1 kb and the data were mined for viral sequences using tools Virfinder (v1.1) and Virsorter v2. Virsorter v1 was used for the identification of viral sequences obtained from the total microbial DNA samples. Extraction of viral contigs with Virfinder was based on a score of >0.7 and p-value < 0.05. Virsorter2 viral contigs were considered with a cut-off score of >0.75. For the bacterial samples, Virsorter-identified contigs belonging to all categories (categories 1-6) were considered for further downstream analysis. All the “non-viral” contigs from Virfinder and Virsorter were taken as input for CAT (Contig Annotation Tool, v5.2). All the viral contigs obtained by the above three tools (VirFinder, VirSorter and CAT) were sorted for each of the 12 samples individually and the collection was run through the CD-HIT-EST (v4.8.1) tool with an identity cut off of 99% over the entire contig length to get non-redundant viral databases for each sample. To analyze the quality of assembled genomes, CheckV (v0.6) tool was used.^30^

For taxonomic identification of the viral contigs, an index was made from Viral RefSeq database 1.1, 2.1 and 3.1 (https://ftp.ncbi.nlm.nih.gov/refseq/release/viral/), using Bowtie2. The Kaiju program was then used to map the contigs to the Viral RefSeq index. For the clustering-based method of taxonomic identification, genes/ORFs were predicted for each of the 12 viral databases using the Prodigal (v2.6.3) tool in metagenomic mode. vConTACT2 was used to cluster and provide a taxonomic context of metagenomic sequencing data. Prodigal GenBank coordinates were used as input for each sample for vConTACT2 and it was run with pc-inflation and vc-inflation set to 1.5, pcs-mode set to MCL, and vcs-mode set to ClusterONE.^31^ Databases used for the analysis were TrEMBL and ProkaryoticViralRefSeq94-Merged.

### 5. crAssphage analysis

To identify crAss-like phages, all assembled metagenomic contigs were BLAST (v2.4.0) searched against a database of crAssphage genomes and proteomes. The database comprised of prototypical crAssphage genome and proteome (p-crAssphage; NC_024711.1), 249 crAssphage genome and 2684 crAssphage family, which were reported by Guerin *et al.,* and Yutin *et al.,* respectively.^23, 27^ From the search result, putative crAss-like phages were selected using the following criteria (i) a BLAST hit against databases with an E-value less than 1E-05, (ii) a BLAST query alignment length 350bp, and (iii) a minimum contig length of 70kb (representing near-complete crAss-like phage contigs).

To determine the phage lifestyle, open reading frames (ORFs) for each crAss-like phage contigs were predicted using Prodigal (v2.6.3) in metagenomic mode (https://github.com/hyattpd/Prodigal). Further, all ORFs were compared to a custom set of 29 HMM profiles that belong to transposase, integrase, excisionase, resolvase and recombinase proteins. The HMM profiles were downloaded from the Pfam database.^32^ Contigs with an ORF, which obtained a hit with any of the above mentioned five functional classes were classified as temperate.^33^

The taxonomic identity of predicted crAss-like phages was done based on average nucleotide identity (ANI) calculated using OrthoANI at default parameters.^34^ ANI of each predicted crAss-like phages were calculated with each of the sixty-five previously identified crAss-like phages genomes classified as genus I to X.^35^ The genus having maximum ANI was assigned as the most probable taxon.

### 6. Determination of species richness and their correlations

Quality filtered reads obtained from the VLP-derived DNA and the total microbial DNA were mapped to a Viral RefSeq index, using ViromeScan2. The index was made from the collection of eukaryotic viruses and bacteriophages present in the Viral RefSeq database using Bowtie2. The output files were mapped to the NCBI taxa accession number using a custom code and an OTU table was generated. Similarly, reads obtained from total microbial DNA were mapped to the bacterial RefSeq database to estimate the abundance of bacterial species, using the MEGAN5 package. A vegan package was applied to calculate the richness of viral and bacterial species in each sample.

### 7. Functional profiles

The non-redundant databases of our viral contigs (>1 kb) from each sample were submitted to the web-server, MG-RAST (**M**etagenomics **R**apid **A**nnotations using **S**ubsystems **T**echnology) (http://rast.nmpdr.org). The parameters used for the assignment to functional categories were e-value 1e-05, % identity cut-off of 60 % and minimum alignment length 15.

### 8. Determination of bacteriophage lifestyle

Viral sequences were clustered into vOTUs using ClusterGenomes v5.1 at 95 % average nucleotide identity (ANI) and 85 % average fraction (AF). ORF in each vOTU were predicted using Prodigal v2.6.3 (https://github.com/hyattpd/Prodigal) in metagenomic mode. All ORFs of each OTU were annotated using a custom set of 29 HMM profiles belonging to transposase, integrase, excisionase, resolvase, and recombinase proteins, using the library of 29 HMM profiles that was originally compiled by Cook et al. from the Pfam database.^33^

## Supplemental Information

Document Supl_Tables

## Acknowledgments

Dr. Bhabatosh Das, Translational Health Science and Technology Institute is gratefully acknowledged for his valuable suggestions on the manuscript.

## Data availability statement

https://dataview.ncbi.nlm.nih.gov/object/PRJNA792685?reviewer=tedv6alaa2mvci3sliad55k7ok

## Author contributions

**KB**: Conception; design; acquisition, analysis and interpretation of data

**HS**: Data collection and final approval

**AG, ADP, MK**: Data analysis, revisiting the manuscript and final approval

**SV**: Conception, revisiting the manuscript and final approval

## Funding

This work was supported by the Department of Biotechnology, Government of India grant no. BT/PR18657/BIC/101/507/2016 to KB. MK acknowledges the grant nos. VIR(25)/2019/ECD-1, and ISRM/12(33)/2019 by the Indian Council of Medical Research, and grant no. EMR/2017/002299 by the Scientific and Engineering Research Board.

## Disclosures

The authors declare no competing interests.

---

**Table S1.** Read Summary.

| Sample i.d. | Number of VLP DNA-derived reads |  | Number of total-microbial DNA-derived reads |  |
| --- | --- | --- | --- | --- |
|  | Total | Mapped to viral RefSeq | Total | Mapped to viral RefSeq |
| **F3** | 55945748 | 27066 ( 0.05 %) | 103310208 | 27083 (0.03 %) |
| **F4** | 68388278 | 60763 (0.09 %) | 117258248 | 21582 (0.02 %) |
| **F5** | 120277474 | 394972 (0.3 %) | 105878448 | 6571 (0.006 %) |
| **F7** | 126813688 | 39176 (0.03 %) | 69697178 | 34207 (0.05 %) |
| **F11** | 26006146 | 38904 (0.15 %) | 106044330 | 66301 (0.06 %) |
| **F12** | 38375058 | 81769 (0.211 %) | 165786894 | 169968 (0.1 %) |
| **F13** | 44020188 | 68081 (0.15 %) | 70267208 | 54502 (0.08 %) |
| **F14** | 38132234 | 26052 ( 0.07 %) | 77844958 | 46839 (0.06 %) |
| **F16** | 50049542 | 41176 (0.08 %) | 100960596 | 36169 (0.04 %) |
| **F18** | 24414020 | 38032 ( 0.16 %) | 42099008 | 29505 (0.07 %) |
| **F21** | 37447036 | 53276 (0.14 %) | 73885912 | 34413 (0.05 %) |
| **F29** | 51299028 | 568553 ( 1.11 %) | 103310208 | 77819 (0.07 %) |
| **Total** | **681168440** | **1437820** | **1136343196** | **604959** |

**Table S2.** Summary of metagenome assembly.

| Sample i.d. | No. of reads |  | No. of total contigs (MEGAHIT) |  | No. of contigs (> 1000 bp) |  | Largest Contig (bp) |  |
| --- | --- | --- | --- | --- | --- | --- | --- | --- |
|  | VLP-DNA | Total Microbial-DNA | VLP-DNA | Total Microbial-DNA | VLP-DNA | Total Microbial-DNA | VLP-DNA | Total Microbial-DNA |
| **F3** | 55945748 | 103310208 | 77253 | 179130 | 11055 | 41248 | 737163 | 725802 |
| **F4** | 68388278 | 117258248 | 48221 | 142785 | 10225 | 38742 | 796189 | 796189 |
| **F5** | 120277474 | 105878448 | 55249 | 454755 | 7655 | 88264 | 640931 | 411643 |
| **F7** | 126813688 | 69697178 | 56594 | 363684 | 7594 | 86724 | 405150 | 330991 |
| **F11** | 26006146 | 106044330 | 15115 | 98940 | 2916 | 24738 | 265523 | 368663 |
| **F12** | 38375058 | 165786894 | 29423 | 109813 | 306 | 21751 | 31161 | 232869 |
| **F13** | 44020188 | 70267208 | 58699 | 174692 | 11561 | 35666 | 176112 | 814305 |
| **F14** | 38132234 | 77844958 | 50041 | 201298 | 7565 | 45857 | 294552 | 1104472 |
| **F16** | 50049542 | 100960596 | 95950 | 360992 | 15590 | 69923 | 535552 | 534186 |
| **F18** | 24414020 | 42099008 | 38350 | 76588 | 5207 | 18440 | 154147 | 537610 |
| **F21** | 37447036 | 73885912 | 34304 | 229747 | 6793 | 55788 | 250352 | 667724 |
| **F29** | 51299028 | 103310208 | 302306 | 259094 | 7145 | 63798 | 489932 | 633385 |
| **Total** | **681168440** | **1239722496** | **861505** | **2651518** | **93612** | **590939** |  |  |

**Table S3.** Summary of data mining to recover viral sequences.

| Sample i.d. | Number of contigs identified as “Viral” |  |  |  |  |  | No. of non-redundant viral contigs |  |  | Length of the recovered viral contigs (bp) |  |  |
| --- | --- | --- | --- | --- | --- | --- | --- | --- | --- | --- | --- | --- |
|  | VirFinder |  | VirSorter |  | CAT |  | VLP-DNA | Total Microbial-DNA | Total | Min-Max | Median | Mean |
|  | VLP-DNA | Total Microbial-DNA | VLP-DNA | Total Microbial-DNA | VLP-DNA | Total Microbial-DNA |  |  |  |  |  |  |
| **F_0_3** | 1576 | 2703 | 1720 | 291 | 68 | 60 | 1905 | 2699 | 4604 | 1000-720556 | 1673 | 6432 |
| **F_0_4** | 1217 | 2184 | 1602 | 354 | 188 | 87 | 1744 | 2230 | 3974 | 1000-584414 | 1825 | 7693 |
| **F_0_5** | 1891 | 5461 | 2376 | 502 | 116 | 46 | 2306 | 5415 | 7721 | 1000-481160 | 1869 | 5919 |
| **F_0_7** | 1915 | 4916 | 2449 | 460 | 298 | 87 | 2944 | 4880 | 7824 | 1000-290467 | 1636 | 5355 |
| **F_0_11** | 665 | 1668 | 771 | 144 | 39 | 33 | 1038 | 1657 | 2695 | 1000-265479 | 1581 | 4739 |
| **F_0_12** | 38 | 1151 | 52 | 143 | 3 | 13 | 35 | 1251 | 1286 | 1000-164209 | 1508 | 4661 |
| **F_0_13** | 1488 | 2709 | 1209 | 249 | 54 | 34 | 2249 | 2628 | 4877 | 1000-289973 | 1745 | 3898 |
| **F_0_14** | 1494 | 3347 | 1122 | 300 | 32 | 12 | 2151 | 3256 | 5407 | 1000-463650 | 1758 | 4655 |
| **F_0_16** | 3004 | 6161 | 2323 | 319 | 55 | 23 | 4071 | 5962 | 10033 | 1000-534202 | 1700 | 4281 |
| **F_0_18** | 834 | 921 | 1034 | 126 | 57 | 4 | 1473 | 961 | 2434 | 1000-431411 | 1601 | 4443 |
| **F_0_21** | 1359 | 4102 | 1039 | 349 | 75 | 41 | 1816 | 4007 | 5823 | 1000-383875 | 1633 | 4466 |
| **F_0_29** | 469 | 3601 | 775 | 432 | 6 | 20 | 642 | 3779 | 4421 | 1000-514315 | 1634 | 5882 |
| **Total** | **15950** | **38924** | **16472** | **3669** | **991** | **460** | **22374** | **38725** | **61099** |  |  |  |

**Table S4.** Summary of CheckV output.

| Sample i.d. | No. of “complete” genomes |  |  | No. of “high quality” genomes (> 90% complete genomes) |  |  | No. of “Undetermined quality” |  |  | No. of “medium quality” (> 50-90% complete genomes) |  |  | No. of “low quality” (> 0-50% complete genomes) |  |  |
| --- | --- | --- | --- | --- | --- | --- | --- | --- | --- | --- | --- | --- | --- | --- | --- |
|  | VLP-DNA | Total Microbial-DNA | Total | VLP-DNA | Total Microbial-DNA | Total | VLP-DNA | Total Microbial-DNA | Total | VLP-DNA | Total Microbial-DNA | Total | VLP-DNA | Total Microbial-DNA | Total |
| F_0_3 | 35 | 12 | 47 | 20 | 16 | 36 | 1221 | 1993 | 3214 | 17 | 24 | 41 | 612 | 654 | 1266 |
| F_0_4 | 51 | 25 | 76 | 21 | 37 | 58 | 1015 | 1596 | 2611 | 31 | 25 | 56 | 637 | 547 | 1184 |
| F_0_5 | 101 | 18 | 119 | 53 | 37 | 90 | 895 | 4164 | 5059 | 60 | 56 | 116 | 1197 | 1140 | 2337 |
| F_0_7 | 79 | 20 | 99 | 37 | 30 | 67 | 1585 | 3733 | 5318 | 42 | 51 | 93 | 1201 | 1046 | 2247 |
| F_0_11 | 13 | 7 | 20 | 14 | 13 | 27 | 639 | 1285 | 1924 | 8 | 13 | 21 | 364 | 339 | 703 |
| F_0_12 | 1 | 2 | 3 | 1 | 8 | 9 | 8 | 878 | 886 | 1 | 6 | 7 | 24 | 357 | 381 |
| F_0_13 | 18 | 9 | 27 | 12 | 18 | 30 | 1527 | 1982 | 3509 | 20 | 12 | 32 | 672 | 607 | 1279 |
| F_0_14 | 17 | 15 | 32 | 10 | 21 | 31 | 1516 | 2494 | 4010 | 11 | 23 | 34 | 597 | 703 | 1300 |
| F_0_16 | 40 | 8 | 48 | 19 | 18 | 37 | 2713 | 4944 | 7657 | 33 | 15 | 58 | 1266 | 977 | 2243 |
| F_0_18 | 21 | 3 | 24 | 10 | 11 | 21 | 590 | 681 | 1271 | 13 | 7 | 20 | 839 | 259 | 1098 |
| F_0_21 | 28 | 21 | 49 | 22 | 18 | 40 | 1236 | 3099 | 4335 | 5 | 23 | 28 | 525 | 846 | 1371 |
| F_0_29 | 33 | 23 | 56 | 14 | 30 | 44 | 273 | 2891 | 3164 | 8 | 33 | 41 | 314 | 802 | 1116 |
| **Total** | **437** | **163** | **600** | **233** | **257** | **490** | **13218** | **29740** | **42958** | **249** | **288** | **547** | **8248** | **8277** | **16525** |

**Table S5.** Identification of crAss-like phages.

| Sample i.d. | Read mapping to crAss-like phages (%) | Number of crAss-like phage contigs (greater than 70 kb; NR) |  |  | Lysogeny-related genes (% crAss-like phages) |
| --- | --- | --- | --- | --- | --- |
|  |  | VLP-DNA | Total Microbial-DNA | Total |  |
| **F3** | 0.1-14 % | 22 | 14 | 36 | 30.5 |
| **F4** | 2-4 % | 19 | 22 | 41 | 39 |
| **F5** | 4-10 % | 19 | 16 | 35 | 40 |
| **F7** | 0.03-1.6 % | 27 | 11 | 38 | 50 |
| **F11** | 0.001 % | 5 | 4 | 9 | 44 |
| **F12** | 0.001 % | 00 | 0 | 0 | NA |
| **F13** | 2-25 % | 4 | 7 | 11 | 36 |
| **F14** | 0-0.2 % | 10 | 4 | 14 | 35 |
| **F16** | 0.04-2 % | 15 | 9 | 24 | 68.8 |
| **F18** | 0-1.8 % | 8 | 6 | 14 | 28.6 |
| **F21** | 0.1-1 % | 10 | 8 | 18 | 61 |
| **F29** | 3-10 % | 5 | 10 | 15 | 26.7 |

